# Impact of neonatal activation of nuclear receptor CAR (Nr1i3) on *Cyp2* gene expression in adult mouse liver

**DOI:** 10.1101/2022.01.21.477223

**Authors:** Aram Shin, David J. Waxman

**Affiliations:** Department of Biology and Bioinformatics Program, Boston University, Boston, MA 02215 USA

**Author notes:** Address correspondence to: Dr. David J. Waxman, Dept. of Biology, Boston University, 5 Cummington Mall, Boston, MA 02115 USA.

**Keywords:** TCPOBOP, phenobarbital, epigenetic reprogramming, growth hormone, Cyp2b10

## Abstract

Perinatal exposure to environmental chemicals is proposed to reprogram development and alter disease susceptibility later in life. Supporting this, neonatal activation of the nuclear receptor CAR (Nr1i3) by TCPOBOP induces persistent expression of mouse hepatic *Cyp2* genes into adulthood, attributed to long-term epigenetic memory of the early life exposure [Hepatology (2012) 56:1499-1509]. Here, we confirm that the same high-dose (15x ED50) neonatal TCPOBOP exposure used in that work induces prolonged (12 weeks) increases in hepatic *Cyp2* expression; however, we show that the persistence of expression can be fully explained by the persistence of residual TCPOBOP in liver tissue. When the long-term presence of TCPOBOP in tissue was eliminated by decreasing the neonatal TCPOBOP dose 22-fold (0.67x ED50), strong neonatal increases in hepatic *Cyp2* expression were still obtained but did not persist into adulthood. Furthermore, the neonatal ED50-range TCPOBOP exposure did not sensitize mice to a subsequent, low-dose TCPOBOP treatment. In contrast, neonatal treatment with phenobarbital, a short half-life (t_1/2_=8 h) agonist of CAR and of PXR (Nr1i2), induced high-level neonatal activation of *Cyp2* genes and also altered their responsiveness to low-dose phenobarbital exposure at adulthood by either increasing (*Cyp2b10*) or decreasing (*Cyp2c55*) expression. Thus, neonatal xenobiotic exposure can reprogram hepatic *Cyp2* genes and alter their responsiveness to exposures later in life. These findings highlight the need to carefully consider xenobiotic dose, half-life and persistence in tissue when evaluating the long-term effects of early life environmental chemical exposures.

## Introduction

Many environmental chemicals dysregulate gene expression, most notably in hepatocytes, by mechanisms that involve activation of members of the Nuclear Receptor superfamily (Toporova and Balaguer 2020; Waxman 1999). CAR and other nuclear receptors are activated by a wide range of structurally diverse foreign chemicals, including many industrial pollutants and pharmaceuticals (Baldwin and Roling 2009; Chang and Waxman 2006; Hernandez et al. 2009; Kobayashi et al. 2015; Omiecinski et al. 2011). CAR also regulates normal physiological pathways, including hepatic energy homeostasis, cell proliferation and inflammation (Cai et al. 2021) and may thereby impact pathophysiological conditions such as fatty liver disease, diabetes, and hepatocellular carcinoma (Cave et al. 2016; Dong et al. 2009; Phillips et al. 2007).

Under normal cellular conditions, CAR is sequestered in the inactive state in the cytoplasm as a multi-protein complex, including heat shock protein 90 and cytoplasmic CAR retention protein (Kobayashi et al. 2003; Timsit and Negishi 2014; Yoshinari et al. 2003). CAR agonist ligands, such as TCPOBOP, bind to the ligand-binding domain of CAR, leading to dissociation of its cytoplasmic chaperones and translocation of CAR to the nucleus, where it heterodimerizes with retinoid X receptor (RXR) (Mackowiak and Wang 2016). The CAR/RXR heterodimer subsequently recruits coactivators and binds to specific response elements in genomic DNA, followed by the transcriptional activation of many genes, including phase I and phase II enzymes of drug metabolism and transporters that regulate the metabolism of endogenous and exogenous chemicals (Mackowiak and Wang 2016; Qatanani and Moore 2005). CAR activation is associated with epigenetic changes in mouse liver proximal to CAR binding sites (Niu et al. 2018; Tian et al. 2018) and nearby CAR responsive genes (Rampersaud et al. 2019). These changes are apparent as early as 3 h after exposure to TCPOBOP and include both increases and decreases in chromatin accessibility, which can be monitored as changes in DNase hypersensitivity (Lodato et al. 2018; Vitobello et al. 2019). Changes in histone methylation and acetylation, and changes in DNA methylation, have also been linked to the activation and repression of CAR target genes in mouse liver (Lempiainen et al. 2011; Rampersaud et al. 2019).

Early developmental exposure to xenobiotics, including chemicals that can activate CAR, has been proposed to lead to neonatal reprogramming (neonatal imprinting) in a way that can alter metabolic function and gene expression in liver and other tissues (Cave 2020; Küblbeck et al. 2020; Piekos et al. 2017). Potential effects on life expectancy (Agrawal and Shapiro 2005) and disease susceptibility later in life have also been reported (Hochberg et al. 2011). The molecular mechanisms that underlie the early developmental lesions that lead to adult pathophysiology are likely to be epigenetic in nature but are poorly understood (Moggs and Terranova 2018; Nahar et al. 2014; Treviño et al. 2020). For example, when CAR is activated in neonatal mouse liver by the CAR-specific agonist TCPOBOP, a persistent increase in expression was reported for two CAR target genes, *Cyp2b10* and *Cyp2c37*, via a proposed epigenetic memory of neonatal CAR exposure (Chen et al. 2012). However, the long half-life apparent for liver metabolic activities induced by TCPOBOP (Poland et al. 1980) raises the question of whether continued presence of TCPOBOP in mouse tissue, rather than an epigenetic memory, drives the observed persistence of CAR target gene induction.

Here, we show that neonatal exposure to TCPOBOP at a dose widely used by many investigators, 3 mg/kg, induces long-term, persistent activation of CAR-responsive genes in the liver, but also results in the persistence of TCPOBOP in liver tissue at a concentration we find is sufficiently high to account for the prolonged increase in expression of *Cyp2* family genes in the liver. Additionally, we characterize the impact of neonatal exposure to phenobarbital, which activates CAR indirectly via changes in CAR phosphorylation (Negishi et al. 2020). Our findings show that neonatal exposure to phenobarbital, which lacks the complications of long-term persistence in tissue owing to its comparatively short half-life (∼8 h) (Markowitz et al. 2010) compared to TCPOBOP, sensitizes mouse liver to a subsequent exposure to phenobarbital in adulthood. Thus, such short-lived chemicals are more amenable for studying mechanisms by which CAR and other xenobiotic sensors translate early environmental stimuli to persistent changes in gene expression.

## Materials and Methods

### Animals

All mouse work was carried out in compliance with procedures approved by the Boston University Institutional Animal Care and Use Committee (protocol # PROTO201800698), and in compliance with ARRIVE 2.0 Essential 10 guidelines (Percie du Sert et al. 2020), including study design, sample size, randomization, experimental animals and procedures, and statistical methods. Adult male and female CD-1 mice (8-10 weeks of age) were purchased from Charles River Laboratories (Wilmington, MA) and housed in the Boston University Laboratory Animal Care Facility. Mice were kept on a 12-hour light cycle (7:30 AM – 7:30 PM). For mouse breeding, one adult male and one adult female mouse were housed together until a vaginal plug was observed, at which time the mice were separated. The first day pups were observed was designated as postnatal day 1 (PND1). Litters were housed with dams until the mice were weaned on PND21. Mice were treated with TCPOBOP (1,4-Bis-[2-(3,5-dichloropyridyloxy)]benzene; Sigma-Aldrich, St. Louis, MO) dissolved in a 1% DMSO solution in tocopherol-stripped corn oil (Fisher Scientific), or with phenobarbital (sodium salt; Sigma-Aldrich) dissolved in 0.9% sodium chloride solution, or with vehicle (control) by intraperitoneal injection between 8:00 AM and 8:45 AM on the day(s) of treatment at doses stated in the text. Treatments were administered to mice at the ages ranging from PND4 to 7 weeks of age, as specified for each study. Mice were euthanized after a time period ranging from 3 h to 51 h after TCPOBOP or vehicle (control) injection, and at a fixed time of day (between 11:00 AM and 11:45 AM) to minimize gene expression variations between mice due to the strong circadian effects on gene expression in liver (Kettner et al. 2016). Where indicated, pregnant dams were given either drinking water supplemented with 0.05% (w/w) phenobarbital or drinking water as a control. A small piece of each liver was snap frozen in liquid nitrogen then stored at -80C prior to extraction of RNA for qPCR analysis and of TCPOBOP for LC/MS analysis.

### Neonatal mouse exposure models

Mice were exposed to either TCPOBOP or phenobarbital during the perinatal period and, where indicated, again later in life, using one of the following four exposure schedules. In Study A (Fig. 1), PND4 pups were given TCPOBOP by i.p. injection at 3 mg/kg or vehicle (control). Mice were given a second injection of TCPOBOP (3 mg/kg) or vehicle in week 3 (on a day between PND20 and PND23) and euthanized 3 h later. In Study B (Fig. 3), PND4 pups were given TCPOBOP at 133 μg/kg (0.67x ED50) or vehicle. Mice were given a second injection of TCPOBOP, at 40 μg/kg (0.2x ED50), or vehicle, in week 7 (on days PND46-PND52) and euthanized 51 h later. In Study C (Fig. 5), newborn mice were exposed to phenobarbital given to nursing dams through their drinking water (0.05% (w/v) phenobarbital) to give a 6 consecutive-day perinatal exposure (PND2 through PND7). In week 7 (on days PND49-PND52), mice were given phenobarbital by i.p. injection at 10 mg/kg/day, or vehicle (control), on each of 3 consecutive days and euthanized 3 h after the last injection. Finally, in Study D (Fig. S2), PND4 pups were given two i.p. injections of phenobarbital, each at 40 mg/kg, or vehicle, on PND4 and again on PND5. In week 7 (on days PND49-50), the mice were injected with phenobarbital at 10 mg/kg/day i.p., or vehicle, on each of 3 consecutive days and euthanized 3 h after the last injection.

**Fig. 1.**
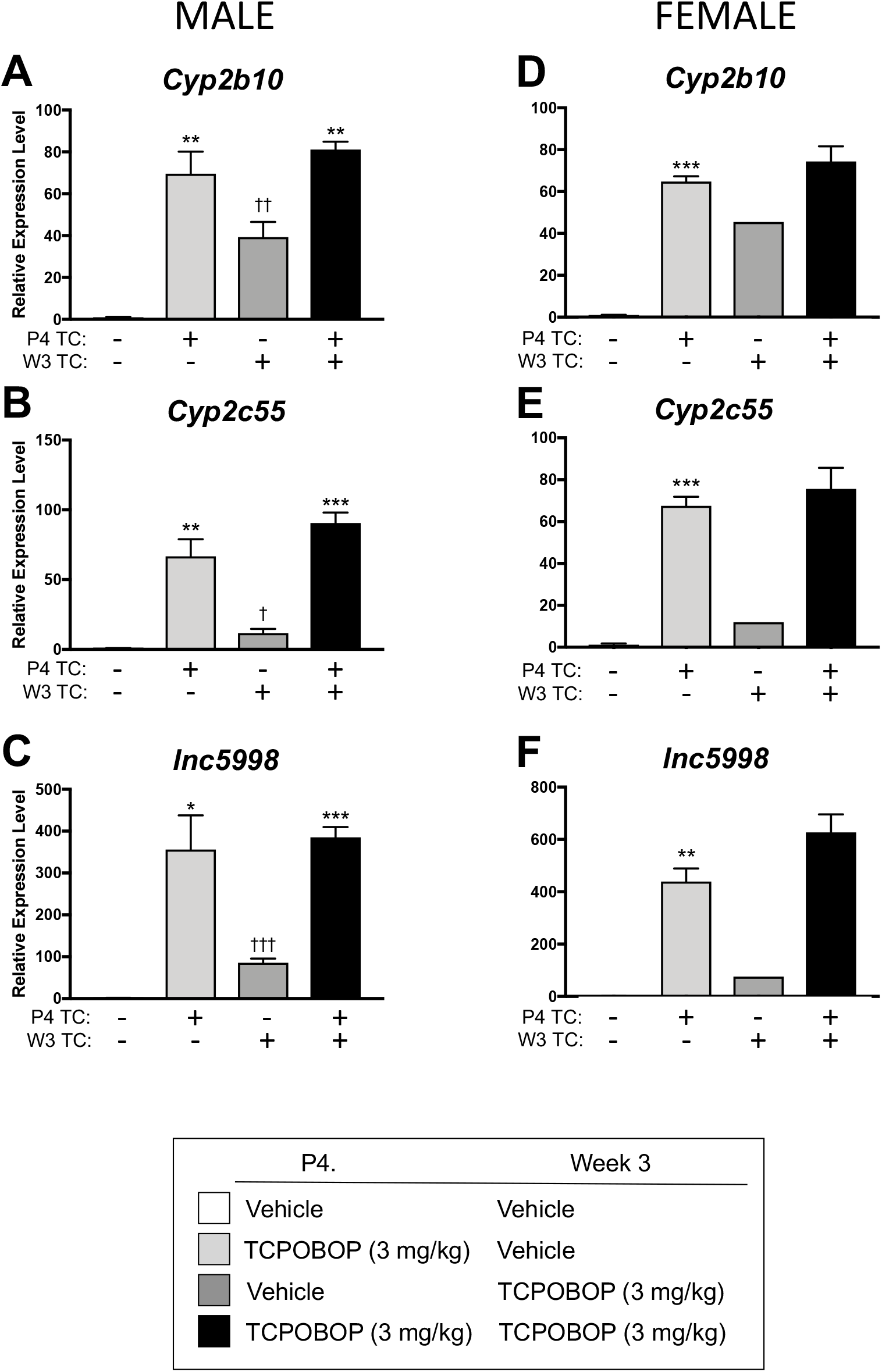
Impact of neonatal TCPOBOP exposure with repeat dosing in week 3 on CAR-responsive genes. Male and female pups were injected with TCPOBOP (3 mg/kg) or vehicle (control) on PND4, and in week 3, were again treated with TCPOBOP (3 mg/kg), or vehicle (control). All mice were euthanized in week 3, 3 h after the final injection (**Study A design**). Shown are relative RNA levels of each gene determined by RT-qPCR analysis of RNA extracted from each liver, with values normalized to that of the vehicle-only group (first bar) set to a value of 1.0. **A-C**, gene expression in male livers; **D-F**, gene expression in female livers. Data are shown as mean +/-SEM (n = 3, 4, 4, 4 individual males, for bars 1-4, respectively, and n = 2, 5, 1, 5 individual females, for bars 1-4, respectively). Data were analyzed by pair-wise t-test for two separate comparisons: *, effects of TCPOBOP exposure on PND4 (bar 2 vs bar 1, and bar 4 vs bar 3); and †, effects of TCPOBOP in week 3 (bar 3 vs bar 1, and bar 4 vs bar 2). TC, TCPOBOP. The same patterns were seen in both sexes, but the small sample size in the week 3 alone female group precluded a full statistical analysis. Significance: * or †, p < 0.05; ** or ††, p < 0.01; *** or †††, p < 0.001.

### RNA purification and RT-qPCR

A portion of each liver (0.1-0.2 g) was homogenized for 30 sec in 1 mL of TRIzol using a Polytron homogenizer. The TRIzol homogenate was processed according to the manufacturer’s protocol (Life Technologies) to purify total liver RNA. The purified RNA was treated with DNase I (Promega) to remove genomic DNA contaminants and then converted to cDNA using Applied Biosystems High Fidelity RT kit (ThermoFisher). Quantitative real time PCR (qPCR) primer pairs specific for mature mRNA were designed to be in adjacent exons of each target gene, with amplicons spanning long introns to reduce the likelihood that amplification of contaminating genomic DNA would contribute to the qPCR signal. Primer pairs specific for primary (unspliced) RNA transcripts were designed to have amplicon span either an exon-injunction or an intron-exon junction of the target gene. qPCR was performed using Power SYBR Green PCR Master Mix (ThermoFisher). Relative RNA expression levels were calculated using the ΔΔCt method and normalized to the expression of 18S ribosomal RNA. Primer sequences used to amplify each gene are shown in Table S1.

### TCPOBOP extraction and LC/MS

A piece of each frozen liver was added to PBS (0.5 g liver/0.5 mL PBS) and immediately homogenized using a Polytron homogenizer for 30 sec. The homogenate (∼ 1 mL) was shaken with 5 mL of hexane for 1 h at room temperature. The hexane layer was transferred to a clean glass tube, and 5 mL of fresh hexane was added to the remaining homogenate and vortexed on a flat platform for 1 h at room temperature. The hexane layers were combined and evaporated under a gentle stream of nitrogen gas. The residue was dissolved in ∼0.1-0.2 mL of 4% DMSO (Sigma-Aldrich) in acetonitrile (Fisher Scientific) and stored at -20C until LC/MS analysis. HPLC was performed using a C18 column run at a flow rate of 0.6 ml/min of 30% water and 70% acetonitrile. The HPLC-separated peaks were further analyzed using a Waters QTof Premier mass spectrometer in the Boston University Chemical Instrumentation Core. Standard curves for TCPOBOP quantification were generated using pure TCPOBOP dissolved in 4% DMSO in acetonitrile. The prominent TCPOBOP total ion count peak at 402.96 m/s was used to calculate the concentration of TCPOBOP present in each liver extract.

### Statistical analysis

Data are presented as mean +/-SEM for the number of individual livers (biological replicates) specified in each figure legend. Significance was assessed by two-tailed t-test for pairwise comparisons specified in each figure legend and implemented in GraphPad Prism.

## Results

### Responsiveness of neonatal mice to TCPOBOP

We investigated the effects of neonatal exposure to TCPOBOP (3 mg/kg on PND4) on liver expression of *Cyp2b10* and *Cyp2c55*, which are both induced >50-fold within 3 h of TCPOBOP treatment in adult mouse liver (Lodato et al. 2017). TCPOBOP stimulated a persistent induction of both Cyp2 genes, as well as of lnc5998, a lncRNA that is highly TCPOBOP-inducible and is divergently transcribed from *Cyp2b10* (Lodato et al. 2017) (Fig. 1, bar 2 vs bar 1). Large, significant inductions were also seen in livers of mice given a single injection of TCPOBOP in week 3 and euthanized 3 h later (bar 3), but the extent of induction was lower than was seen in the PND4-treated mice (bar 3 vs bar 2) or in mice exposed to TCPOBOP on both PND4 and at 3 weeks of age (bar 3 vs bar 4). Similar results were observed in female mice (Fig. 1A-C vs Fig. 1D-F). The increased effectiveness of TCPOBOP at inducing gene expression in mice receiving both TCPOBOP treatments was much greater for *Cyp2c55* and *lnc5998* than for *Cyp2b10*, which both respond to a single injection of TCPOBOP more slowly than *Cyp2b10* (Lodato et al. 2017), and hence are not maximally induced at the 3 h time point.

Although all three genes were significantly induced by both neonatal and week 3 TCPOBOP exposure, we did not observe a further increase beyond that induced by neonatal TCPOBOP in either sex when the mice were challenged with a second injection of TCPOBOP at 3 weeks of age (Fig. 1, bar 4 vs bar 2). Given the long half-life of TCPOBOP, about 2 weeks in adult mice (Poland et al. 1980), these findings suggest that the transcriptional activation of these CAR target genes reaches its maximal level following the first TCPOBOP injection on PND4, and that the higher expression at week 3 seen in PND4-treated mice compared to week 3 (3 h)-treated mice (bar 2 vs. bar 3) reflects the much longer effective exposure time in the PND4 treatment group (17 days vs 3 h).

#### Optimization of neonatal and TCPOBOP dose

We sought to distinguish any potential long-term reprogramming effects of neonatal TCPOBOP exposure from effects due to the residual TCPOBOP that may persist in liver or other tissues. We first optimized the dose of TCPOBOP used in the initial, neonatal exposure. We reasoned that the dose needs to be high enough to activate a robust liver gene response but sufficiently low to be effectively cleared within a few weeks, i.e., to a level low enough to discern the impact of a second TCPOBOP exposure later in life. PND4 mice were given TCPOBOP at doses of 0.33x, 0.67x and 1x the ED50 dose, based on the ED50 value of 0.2 mg TCPOBOP/kg body weight reported for induction of hepatic cytochrome P450-dependent aminopyrine N-demethylase activity in 6 week female mouse liver (Poland et al. 1980). Of note, the ED50 value of 0.2 mg/kg is 15-fold lower than the standard dose of 3 mg/kg TCPOBOP that is widely used in mouse liver studies of CAR target gene induction, including Fig. 1 and in (Chen et al. 2012). Fig. 2 shows that the expression of *Cyp2b10* and *Cyp2c55* was detectable but minimally responsive to an ED50 dose of TCPOBOP after 3 h and then increased dramatically by the 27 h time point (Fig. 2A, Fig. 2B). Given the rapid induction of both genes within 3 h when mice are given the saturating dose of 15x ED50 (i.e., 3 mg/kg) (Lodato et al, 2017), we surmise that the slower induction time course seen with ED50-range doses of TCPOBOP (Fig. 2) reflects the relatively long time required for TCPOBOP to biodistribute to the liver and generate a tissue level sufficient to meet the threshold concentration for strong activation of CAR and its target genes. Very similar patterns were seen for the dose-response and time course of induction of the primary, unspliced transcripts *Cyp2b10* and *Cyp2c55* (Fig. 2C, Fig. 2D), which are indicative of relative rates of gene transcription due to the expected short half-life of such transcripts (Gaidatzis et al. 2015). The apparent increases in *Cyp2* gene transcription rates from 3 h to 27 h, as well as the TCPOBOP dose-dependent increases seen at 27 h, are consistent with an increase in the abundance of transcriptionally active hepatic CAR−TCPOBOP complexes from 3 h to 27 h.

**Fig. 2.**
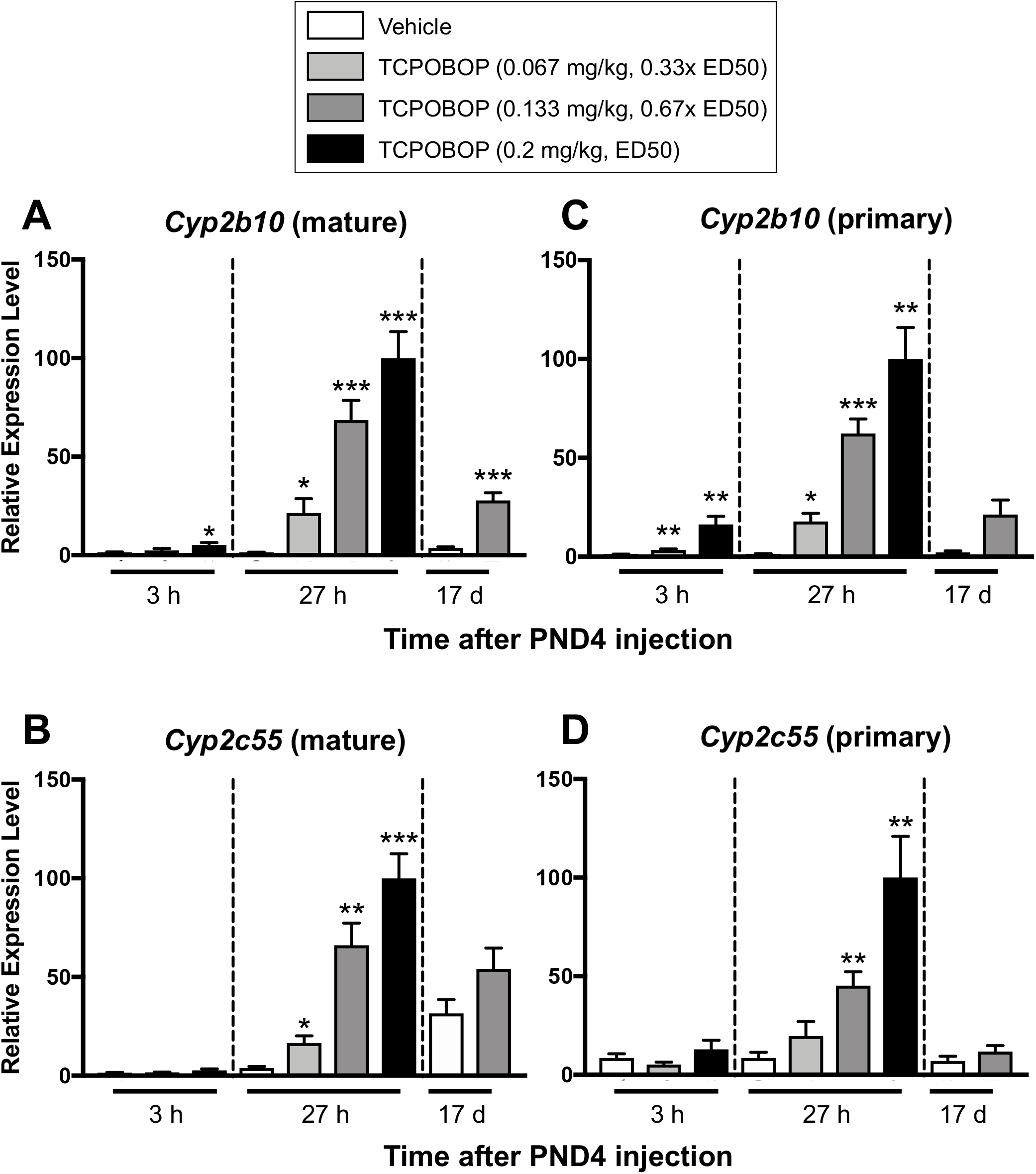
TCPOBOP dose-response in neonatal mouse liver. Male and female PND4 pups were injected with TCPOBOP at 0, 66.7 μg/kg (0.33x ED50), 133 μg/kg (0.67x ED50), or 200 μg/kg (ED50) and euthanized after 3 h (on PND4), after 27 h (on PND5), or after 17 d (on PND21). Shown are gene expression data for the mature and primary RNA transcripts of *Cyp2b10* and *Cyp2c55* for n = 4 to n = 8 livers per group. Data presentation as in Fig. 1, with each experimental condition compared to its age-matched control (first bar at each time point): *, p < 0.05; **, p < 0.01; and ***, p < 0.001. All Y-axis values are relative to 0.2 mg/kg TCPOBOP, which was set = 100.

The induction of Cyp2b10 mature mRNA and its transcription rate (i.e., primary transcript level) 27 h after TCPOBOP exposure on PND4 showed a strong decrease 17 d later, on PND21 (Fig. 2A, Fig. 2C). The magnitude of this decrease is consistent with the 14-day half-life for TCPOBOP elimination from the liver (Poland et al. 1980) when taking into account the substantial increase in body size and liver weight from PND4 to PND21, which will effectively dilute the residual TCPOBOP concentration in liver. Thus, 17 days after TCPOBOP injection on PND4 at a dose of 0.67x ED50 (0.133 mg/kg), there is sufficient elimination of TCPOBOP to decrease Cyp2b10 expression significantly. The transcriptional rate (primary transcript level) of *Cyp2c55* was also low after 17 days compared to 27 h. In contrast, mature Cyp2c55 mRNA levels were elevated after 17 days (i.e., on PND21), both with and without TCPOBOP, indicating there is a developmental accumulation in the basal level of Cyp2c55 mRNA, a general characteristic of many genes during this period of liver maturation (Gunewardena et al. 2015). Based on these findings, we selected an 0.67x ED50 dose of TCPOBOP (0.133 mg/kg) to evaluate the potential of neonatal TCPOBOP exposure for reprogramming later in life.

#### Optimization of adult challenge TCPOBOP dose

Next, we sought to identify a suitable dose of TCPOBOP to use for a subsequent exposure, when adult mice exposed to TCPOBOP neonatally are challenged with a second TCPOBOP injection. We reasoned that the second, challenge dose of TCPOBOP needs to induce CAR-responsive genes significantly, but to a level that is less than maximal, which would allow us to detect any additive or synergistic gene induction due to the impact of the prior, neonatal exposure. Male mice, 7-weeks of age, were treated with TCPOBOP at 0.05x, 0.2x, and 1x ED50 doses and livers were harvested two days later after 51 h. This time point was chosen to give sufficient time for TCPOBOP to biodistribute to the liver and induce gene expression. Fig. S1 shows that TCPOBOP at a 0.2x ED50 dose (0.04 mg/kg) increased *Cyp2* gene expression significantly, but to a level that was sub-maximal; thus, gene responses were 5.4 to 5.6-fold lower (Cyp2b10) or 3.4 to 8.5-fold lower (Cyp2c55) than at the 1x ED50 dose for both mature and primary gene transcripts. Non-linear dose-responses were apparent when comparing the 0.05x and 0.2x ED50 doses, which could be due to the retention in fat (Poland et al. 1980) of a higher fraction of the administered TCPOBOP dose when mice are treated at the lowest dose.

#### Combination of neonatal TCPOBOP with adult rechallenge exposure

Based on the above dose-response studies, we injected mice with TCPOBOP at 0.67x ED50 on PND4 and then re-challenged the mice with a second TCPOBOP exposure at 0.2x ED50 in week 7. Livers were harvested 51 h later and analyzed for expression of CAR-responsive genes. PND4 TCPOBOP treatment alone had no discernable effect on *Cyp2b10* or *Cyp2c55* expression at 7 weeks, in either males or females (Fig. 3, bar 2 vs bar 1). The low, sub-maximally inducing re-challenge dose of TCPOBOP given in week 7 had the expected short-term inductive effect on CAR target gene expression (Fig. 3, bar 3 vs bar 1), but did not result in any additive or synergistic response when given to mice treated with TCPOBOP neonatally (Fig. 3, bar 4 vs bar 3). Thus, neonatal exposure to TCPOBOP does not lead to persistent induction of these CAR target genes, nor does it sensitize these genes to a subsequent exposure at adulthood in either sex.

**Fig. 3.**
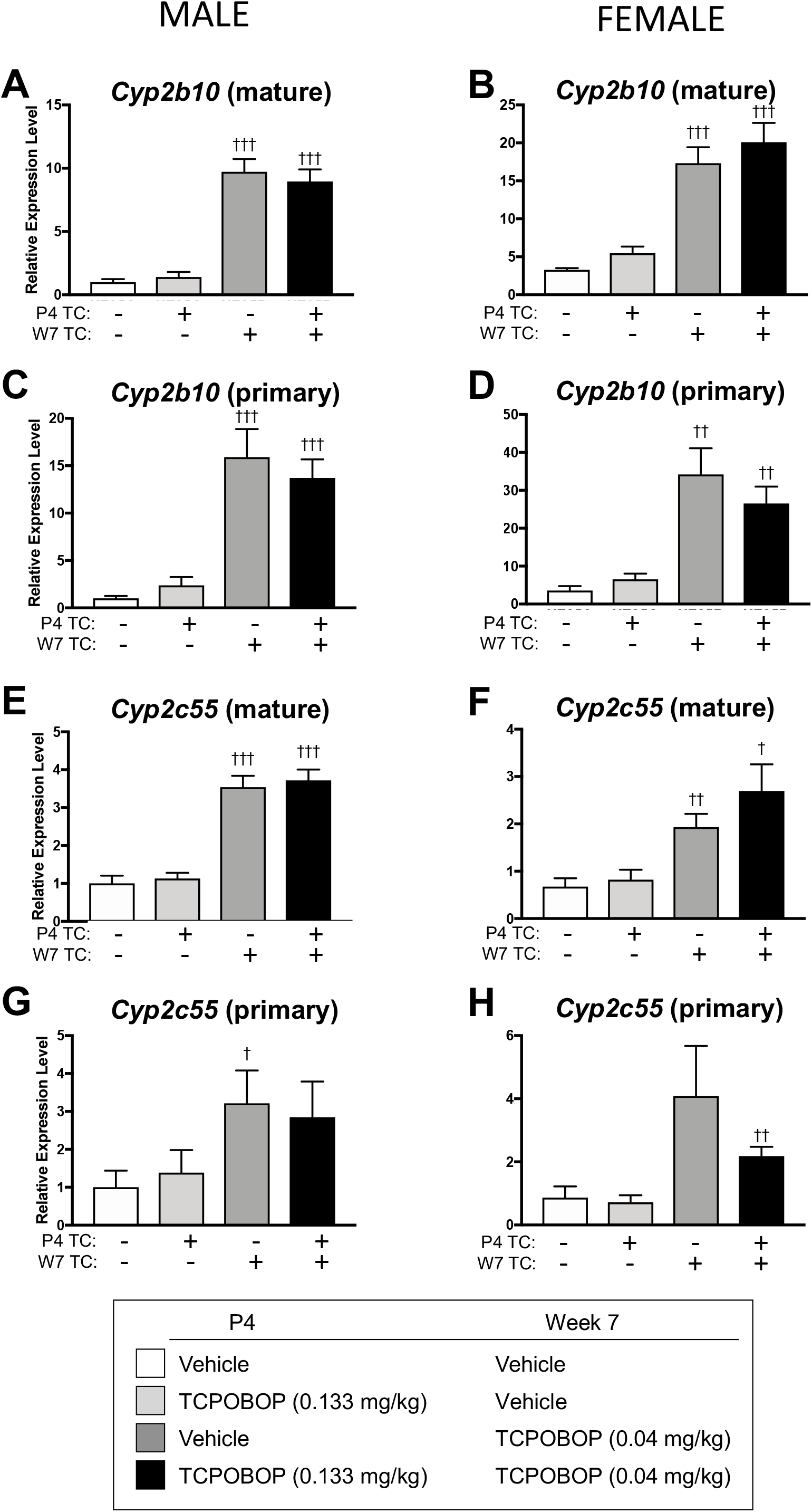
Neonatal TCPOBOP exposure followed by second TCPOBOP exposure in week 7. Male and female PND4 pups were injected with TCPOBOP at 133 μg/kg (0.67x ED50), or vehicle (control), and in week 7 were re-challenged with a second dose of TCPOBOP at 40 μg/kg (0.2x ED50) or with vehicle (control). All mice were euthanized in week 7, 51 h after the last injection (**Study design B**). Shown are the expression levels of the mature and primary transcripts of *Cyp2b10* and *Cyp2c55*, as indicated, for n = 5-7 livers per group. Data presentation as in Fig. 1, comparing the effects of TCPOBOP exposure in week 7 alone (bar 3 vs bar 1) or in combination with TCPOBOP exposure on PND4 (bar 4 vs bar 2): †, p < 0.05; ††, p < 0.01; and †††, p < 0.001. The effects of TCPOBOP exposure on PND4 were not significant by week7 for any of the genes (bar 2 vs bar 1, and bar 4 vs bar 3). Y-axis values are expressed relative to the vehicle-treated control male group for each gene.

Given the absence of any reprogramming effects of neonatal TCPOBOP exposure when mice were exposed to low dose (0.67x ED50) TCPOBOP neonatally (Fig. 3), we sought to confirm the persistent induction of *Cyp2* family genes that was previously seen at week 12 when neonatal mice were treated with TCPOBOP at a high dose, 15x ED50 (3 mg/kg) (Chen et al. 2012). Male and female PND4 pups were treated with TCPOBOP (3 mg/kg), and livers were collected at weeks 3, 7 and 12. Gene expression levels were compared to those seen in livers of week 7 male mice treated with a range of TCPOBOP doses (0, 0.2x, 1x, and 15x ED50) and euthanized 51 h later (Fig. 4A). Neonatal TCPOBOP exposure at 15x ED50 induced persistent expression of *Cyp2b10* and *Cyp2c55* after both 7 and 12 weeks at levels significantly higher than the control group (Fig. 4B and Fig. 4C, groups G-J vs A). Moreover, the expression levels at week 7 were very similar to those seen in livers of 7-week mice exposed to low dose TCPOBOP (0.2x ED50) for 51 h (group B). Next, we employed LC/MS analysis to measure hepatic TCPOBOP concentrations to determine whether the persistent expression of these genes can be explained by TCPOBOP remaining in the liver at 7 weeks. Fig. 4D shows that residual TCPOBOP persists in liver tissue at week 7 and at week 12 at a level similar to the level found 51-h after TCPOBOP dosing at 0.2x ED50 (groups G-J vs A), which is also the neonatal dose that induces an equivalent level of persistent *Cyp2* expression (i.e., G-J vs A in Fig. 4B, Fig. 4C). Thus, the persistent expression of both *Cyp2* genes under the conditions of TCPOBOP treatment used here and by others (Chen et al. 2012) can be fully explained by the continued presence of TCPOBOP in liver tissue even 12 weeks after the initial dosing.

**Fig. 4.**
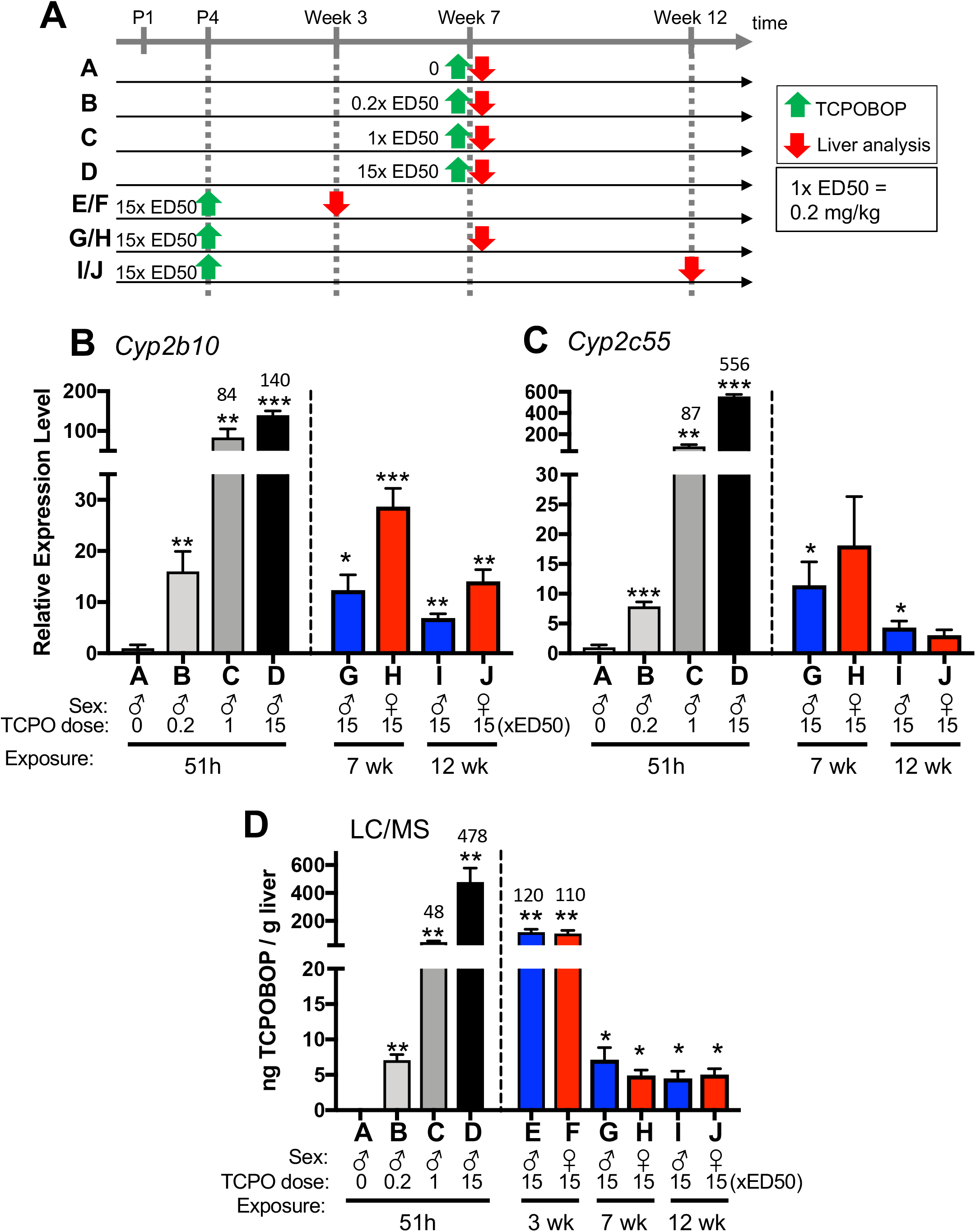
Neonatal exposure to TCPOBOP (3 mg/kg) results in significant residual TCPOBOP and persistent *Cyp2* gene expression after 7 and 12 weeks. **A**: Time course and experimental design. Green arrows, TCPOBOP treatment; red arrows, collection of liver for analysis. Male and female PND4 pups were injected with 3 mg/kg of TCPOBOP (15x ED50 dose) and were euthanized in week 3, 7 or 12 (groups E-J, green and red arrows). For comparison, 7-week-old male mice were injected with TCPOBOP at 0, 0.2x, 1x, and 15x ED50 doses (0, 0.04, 0.2, or 3 mg/kg, respectively) and were euthanized 51 h later (groups A-D; gray and black bars in panels B-D). Further details about each group are shown along the x-axis of panels B-D. **B, C**: RT-qPCR analysis of liver RNA, with expression levels of *Cyp2b10* and *Cyp2c55* normalized to group A (vehicle control). **D**: TCPOBOP was extracted from 0.5 g of each liver with hexane and quantified by LC/MS. In B-D, data shown are mean and SEM, n = 4 per group. Values above the tallest bars indicate the actual y-axis values. Significance compared to vehicle control (group A): *, p < 0.05; **, p < 0.01; ***, p < 0.001.

#### Long-term effects of neonatal phenobarbital exposure

Taken together the studies above demonstrate that: 1) neonatal exposure to a low dose TCPOBOP (0.67x ED50) does not lead to persistent *Cyp2* gene expression at week 7; and 2) neonatal exposure to a high dose of TCPOBOP (15x ED50) does not allow us to evaluate its potential reprogramming effects later in life due to the persistent elevation of TCPOBOP levels in the liver. It remains possible, however, that *Cyp2* genes can be reprogrammed for persistent expression, but that this requires a high level of CAR activation in neonatal liver. We tested this hypothesis in neonatal mice exposed to phenobarbital, a CAR agonist with a much shorter half-life than TCPOBOP (t_1/2_ (phenobarbital) = 15.8 h in PND19 mice (CD-1 strain), and t_1/2_ = 7.5 h in adult mice (NMRI strain) (Markowitz et al. 2010).

For this study, we widened the window of neonatal phenobarbital exposure to encompass days PND2 through PND7 to include critical early time periods that may be required for gene reprogramming, as has been reported for several chemical exposures (Hanson and Gluckman 2014; Hanson and Skinner 2016; Vickers 2011). Phenobarbital was delivered via the drinking water consumed by the dams, an established route for perinatal exposures (Waalkes et al. 2003) that enables drug delivery to neonatal mice by lactation (Asoh et al. 1999). Pups were weaned on PND21 and then left untreated until a second, challenge exposure to phenobarbital was given by i.p. injection at 7 weeks of age. We found that early phenobarbital exposure from PND2-PND7 did not lead to persistent expression of *Cyp2b10* or *Cyp2c55* at week 7 (Fig. 5, bar 2 vs bar 1). Further, in females, we observed a moderate but significant increase in expression of the mature *Cyp2b10* transcript after phenobarbital re-challenge in week 7. The same trend was seen in males for the mature *Cyp2b10* transcript, and in both sexes for the primary *Cyp2b10* transcript, but without reaching statistical significance (Fig. 5A-5D). In contrast, *Cyp2c55* showed a significant decrease in expression in female but not male liver after the phenobarbital rechallenge, as was seen for both the mature and the primary transcripts.

**Fig. 5.**
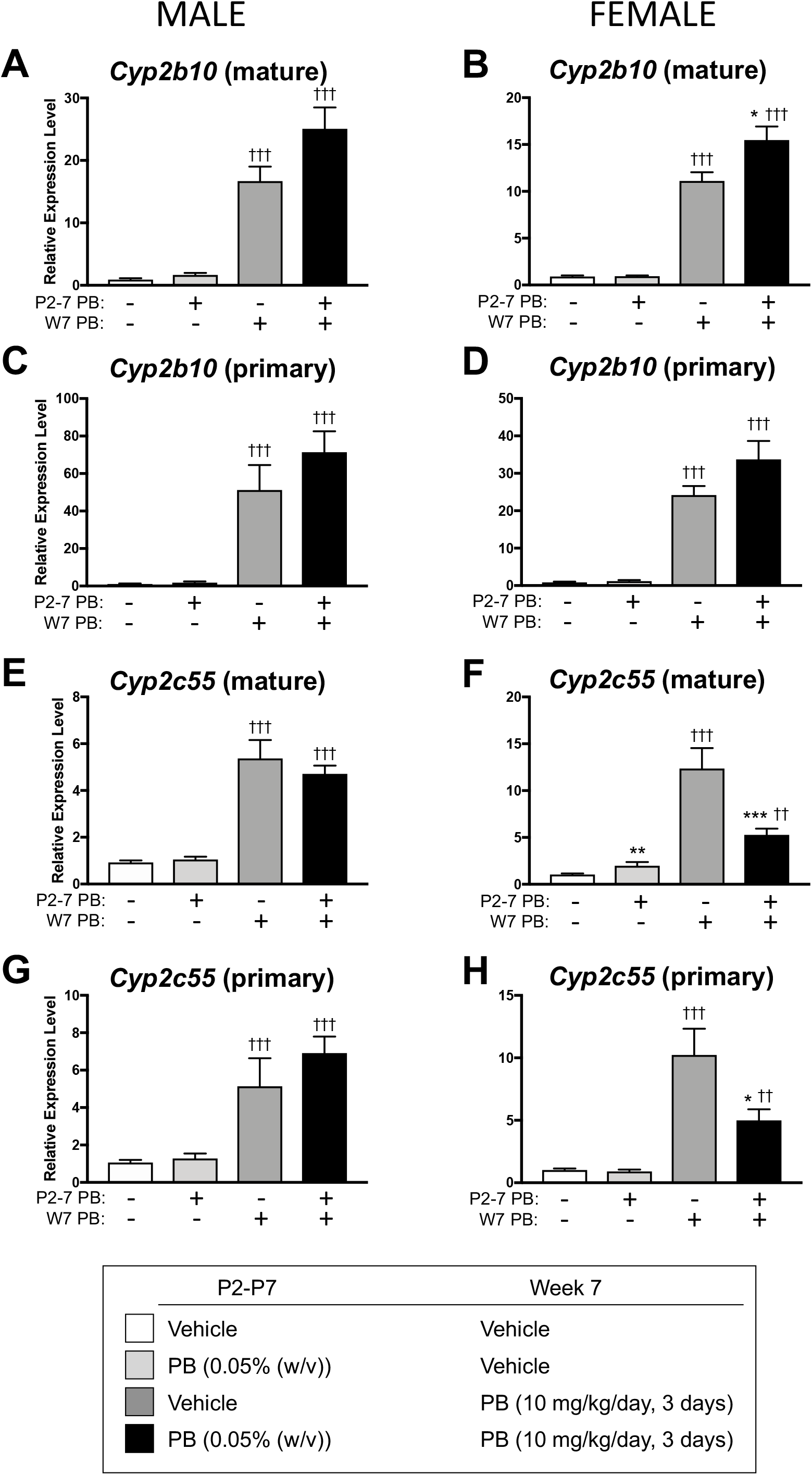
Neonatal phenobarbital exposure (0.05% (w/v) in drinking water fed to the dams) from PND2 to PND7, with a second phenobarbital exposure (10 mg/kg, daily for 3 days), or vehicle control, given in week 7. All mice were euthanized in week 7, 3 h after the last injection (**Study design C**). Shown are RT-qPCR expression data for mature and primary transcripts for *Cyp2b10* and *Cyp2c55*, normalized to the vehicle control (first bar in each set), mean +/-SEM (n = 21, 12, 5, 12 individual males, for bars 1-4, respectively, and n = 23, 8, 8, 14 individual females, for bars 1-4, respectively). Data presentation as in Fig. 1, comparing the effects of phenobarbital exposure in week 7 alone (bar 3 vs bar 1) or in week 7 in combination with PND2-PND7 phenobarbital exposure (bar 4 vs bar 2): *, comparison of bar 2 vs bar 1, and of bar 4 vs bar 3; and †, comparison of bar 3 vs bar 1, and of bar 4 vs bar 2. Significance: * or †, p < 0.05; ** or ††, p < 0.01; *** or †††, p < 0.001.

These results were largely confirmed in a second study, where neonatal mice were injected with phenobarbital at 40 mg/kg/day on days PND4 and PND5, and then re-challenged with phenobarbital at 10 mg/kg/day on 3 consecutive days in Week 7 (Fig. S2). Again, neonatal phenobarbital exposure alone did not lead to persistent expression after 7 weeks, but when combined with adult phenobarbital exposure in week 7, increased the level of *Cyp2b10* mature transcript in male mice significantly higher than that seen in mice given the week 7 exposure alone (Fig. S2, bar 4 vs bar 3). In females but not males, *Cyp2c55* showed a decreased response to the adult phenobarbital re-challenge, very similar to the decrease seen in Fig. 5F and Fig. 5H, but this effect did not reach statistical significance. We conclude that neonatal exposure to a high dose of phenobarbital induces a moderate reprogramming of these *Cyp2* genes in week 7. These reprogramming effects of early phenobarbital exposure are not indirect responses to changes in CAR expression, whose expression did not change significantly with these treatments (Fig. S3A).

#### Impact on sex-specific liver gene expression

Perinatal exposure of rats to phenobarbital can cause long-term dysfunctions such as changes in circulating hormone levels, decreased levels of hepatic monoamine oxidase and infertility (Agrawal et al. 1995; Gupta et al. 1982; Soliman and Richardson 1983). In particular, neonatal phenobarbital exposure was found to alter pituitary secretion patterns and circulating blood levels of growth hormone later in life, and correspondingly, it alters adult liver expression patterns of *Cyp* genes that show sex-dependent, growth hormone-regulated expression patterns (Agrawal and Shapiro 2003). Here, we investigated the effects of both neonatal and 7-week phenobarbital exposure on the adult liver expression of two sex-specific, growth hormone-regulated genes, the female-specific *A1bg* and the male-specific *Cyp7b1* (Fig. S5). Neonatal phenobarbital exposure had no effect on the expression of either gene at 7 weeks of age. Furthermore, adult exposure to phenobarbital, at 7 weeks of age, led to only a small increase in *Cyp7b1* expression. These findings suggest that in the mouse model, early phenobarbital exposure does not lead to major changes in plasma growth hormone profiles known to regulate these sex-specific genes. These findings, in turn, suggest that the effects of neonatal phenobarbital on *Cyp2* genes, described above, are unlikely to be an indirect response to circulating growth hormone levels.

## Discussion

Exposure to endocrine-active environmental chemicals during the perinatal period, a critical window of developmental plasticity, has been proposed to reprogram development and alter disease susceptibility later in life. Many environmental chemicals disrupt gene expression by interaction with xenobiotic sensors from the Nuclear Receptor superfamily, including CAR, which coordinates cellular and transcriptional responses affecting hepatic drug metabolism, energy homeostasis and tumor development. A prior study found that TCPOBOP injection in PND4 mice induced long-term hepatic expression of several CAR target genes from the *Cyp2* family. Elevated expression persisted into adulthood in association with long-term epigenetic changes (Chen et al. 2012), which could involve activation of an epigenetic switch (Lempiainen et al. 2011). However, given the long biological half-life of TCPOBOP (t_1/2_ ∼14 days) (Poland et al. 1980), we asked whether persistence of TCPOBOP in mouse tissue, rather than an epigenetic memory, might drive the persistence of gene expression. We confirmed that the neonatal TCPOBOP exposure regimen used by (Chen et al. 2012) does indeed induce long-term increases in liver *Cyp2* expression lasting at least 12 weeks; however, the persistence of expression was readily explained by the persistence of TCPOBOP in liver tissue at a level sufficient to account for the prolonged increase in expression that we observed. We were able to avoid the long-term persistence of TCPOBOP in liver tissue by decreasing the neonatal TCPOBOP exposure dose 22-fold, from a dose of 15x ED50 used in (Chen et al. 2012)) to a dose of 0.67x ED50. However, although strong neonatal increases in hepatic *Cyp2* expression were still achieved, they did not persist into adulthood. Moreover, this early ED50-range exposure to TCPOBOP did not sensitize mice to a subsequent, low-dose TCPOBOP dosing. Thus, tissue persistence of TCPOBOP, rather than an epigenetic memory, drives the persistence of the elevated *Cyp2* gene expression seen in mouse liver. These findings highlight the importance of carefully considering both dose and pharmacokinetics of elimination from tissue depots when evaluating chemicals such as TCPOBOP for potential reprogramming effects of early life exposures.

Persistent local epigenetic changes were also reported by (Chen et al. 2012) in the neonatal TCPOBOP exposure model; however, those same epigenetic changes are also induced by short-term TCPOBOP exposure (Rampersaud et al. 2019). Consequently, we can attribute them to the ongoing activation of CAR by residual TCPOBOP in liver tissue, rather than to a long-term epigenetic memory of the initial exposure. TCPOBOP has also been shown to accumulate in mouse maternal adipose tissue, from where it can be transferred to pups by lactation to activate CAR-responsive genes in offspring livers (Dietrich et al. 2018). Exposures via that route may appear to give rise to transgenerational effects on gene expression or epigenetics, when in fact they are directly linked to the parental exposure. For TCPOBOP and other lipophilic chemicals, it can thus be difficult to distinguish true long-term gene dysregulation from long-term changes due to ongoing exposure via tissue depots that persist in liver, fat or elsewhere. For such chemicals, reducing the exposure dose to effectively shorten the overall exposure period, as was done here for TCPOBOP, is one approach to determine a chemical’s intrinsic potential for persistent biological effects, albeit with the caveat that in some cases a higher dose may be needed to elicit a robust, long-term epigenetic response.

Accordingly, we used phenobarbital to test whether a threshold level of CAR activation, perhaps not reached by the 0.67x ED50 dose of TCPOBOP, may be required for long-term *Cyp2* reprogramming. This short half-life CAR agonist (t_1/2_ ∼8 hr in adult mice) enabled us to achieve high level neonatal CAR activation without the persistent exposure that is unavoidable when using correspondingly high doses of TCPOBOP due to its ∼50-fold longer half-life. Neonatal phenobarbital induced long-term changes in the responsiveness of *Cyp2* genes to a second, low dose exposure to phenobarbital at adulthood. Specifically, we observed moderate increases in the responsiveness of *Cyp2b10* to low dose phenobarbital at week 7, as well as decreased responsiveness of *Cyp2c55* in female but not male liver, as was seen in two neonatal exposure models with different designs. These findings support earlier work in the rat model, where neonatal phenobarbital administration led to 30-40% over-induction of rat hepatic *CYP2B1* and *CYP2B2* expression when the rats were rechallenged as adults with low doses of the barbiturate (Agrawal and Shapiro 1996). Persistent induction of mouse hepatic CAR target genes was previously described following neonatal phenobarbital exposure (Tien et al. 2015), however, we found that the LD50-range dose of phenobarbital used in that study (>200 mg/kg on PND5) was severely toxic with some lethality (unpublished experiments) and is thus not pharmacologically relevant. Further investigation will be required to elucidate underlying mechanisms, including why the early life exposure to phenobarbital employed in our study has opposite effects on these *Cyp2b10* and *Cyp2c55*, both of which are themselves strongly induced by the initial phenobarbital treatment. Moreover, as phenobarbital activates both CAR and the related nuclear receptor PXR, and with significant overlap between their target genes (Cui and Klaassen 2016), it will be important to determine which receptor mediates the long-term gene responses to neonatal phenobarbital exposure described here.

We initially selected TCPOBOP for studying long-term effects of early CAR activation due to its high specificity for a single nuclear receptor, CAR (Tojima et al. 2012; Tzameli et al. 2000), and for the unusually strong gene responses it can induce, as exemplified by the *Cyp2* genes examined here. While TCPOBOP is a specific activating ligand of rodent but not human CAR owing to species-specific differences in CAR’s ligand binding domain (Mackowiak and Wang 2016), DNA-binding and the associated key genomic and epigenetic effects of CAR activation are most likely conserved across mammalian species. Studies of indirect CAR agonists, such as those reported here for phenobarbital, may be more readily extrapolated across species due to conservation of the overall signaling pathways through which they activate CAR. Phenobarbital activates CAR by inhibiting epidermal growth factor receptor signaling (Chai et al. 2016), which ultimately leads to dephosphorylation of CAR-threonine 38 and CAR nuclear translocation (Mackowiak and Wang 2016; Qatanani and Moore 2005). One limitation, however, is that multiple receptors are often activated by xenobiotics that dysregulate gene expression in the liver (i.e., both CAR and PXR in the case of phenobarbital), which as noted complicates mechanistic studies of any downstream epigenetic actions. Epigenetic reprogramming may also occur in adult mouse exposure models, where long-term treatment with phenobarbital induces many novel differentially methylated and hydroxymethylated genomic regions that strongly correlate with transcriptional responses and are not found after a short-term exposure (Thomson et al. 2013).

Finally, our findings have implications for studies such as those of the Target II consortium (Wang et al. 2018), where multiple environmental chemical exposures are being investigated in perinatal mouse models, including evaluation of changes in gene expression, changes in chromatin accessibility and other epigenetic changes at adulthood. Understanding the mechanisms underlying these processes will provide important insight into how exposure to environmental chemicals and pharmaceuticals in early life can influence long-term health outcomes and disease risk. Challenges going forward will include distinguishing long vs short term effects for chemicals with long half-lives, determining which receptors mediate the effects observed and elucidation of underlying mechanisms, including mechanisms driving epigenetic changes that are expected to be a main driver of long-term phenotypes seen following many environmental chemical exposures.

## Authors’ contributions

AS and DJW jointly conceived of the study. All of the experimental work, data analysis and figure preparation was performed by AS. DJW drafted the manuscript with input and review by AS. DJW provided guidance and supervised the overall project and revised and finalized the manuscript for publication. Both authors reviewed and approved the final manuscript.

## Acknowledgements

The authors thank Dr. Norman Lee, Dept. of Chemistry, Boston University, for assistance with mass spectrometry analysis.

## Supplementary Figures

**Fig. S1.**
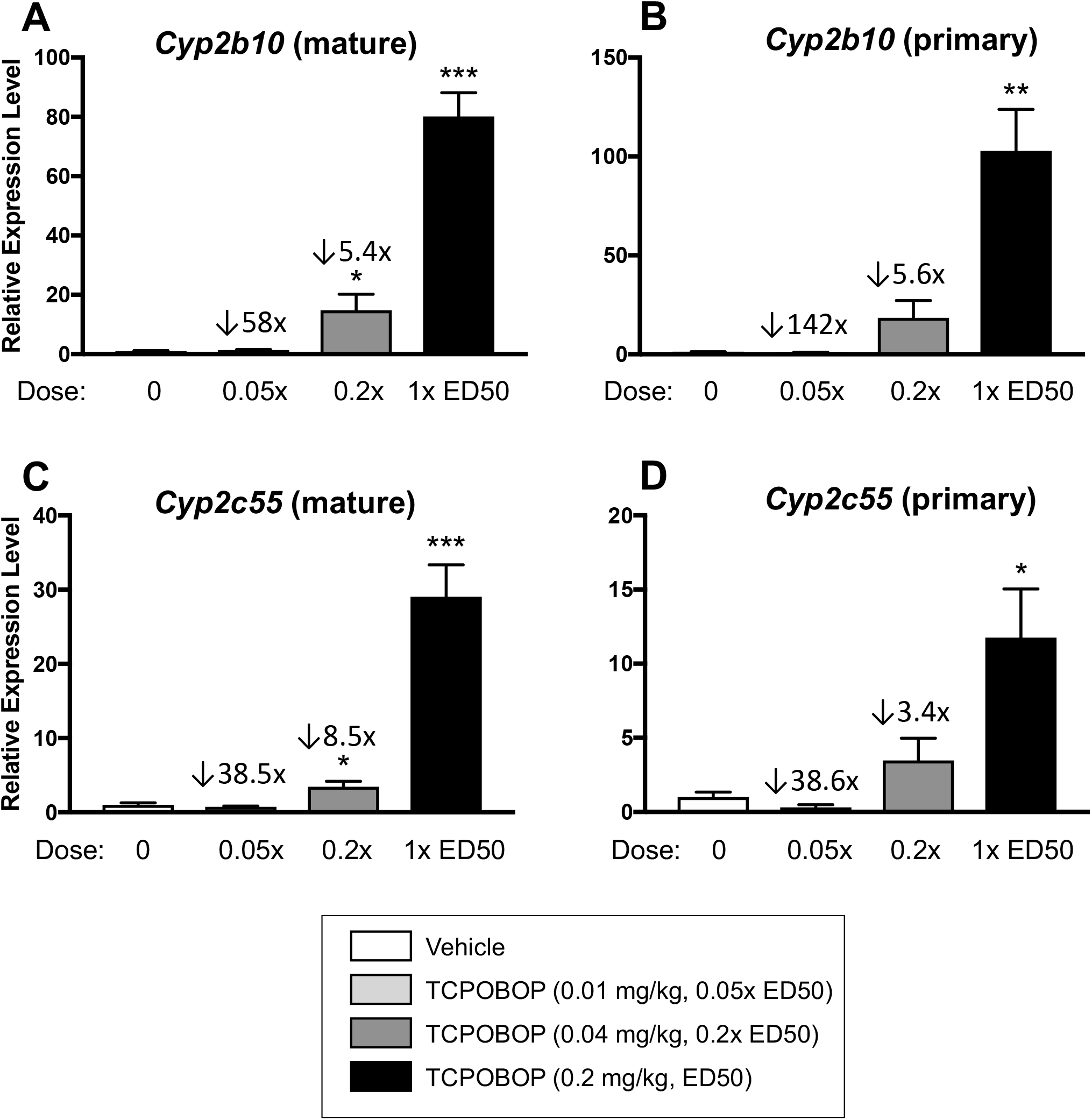
TCPOBOP dose-response in 7-week-old male mice. Mice were injected with TCPOBOP intraperitoneally at 0, 10 μg/kg (0.05x ED50), 40 μg/kg (0.2x ED50), or 200 μg/kg (1x ED50), and euthanized at 51 h later. Total liver RNA was extracted and analyzed by RT-qPCR. Expression levels were normalized to that of the vehicle control group. Shown are the expression levels of the mature and primary transcripts of Cyp2b10 and Cyp2c55, mean and SEM error bar for n = 5 livers per group. Values above the 0.05x ED50 and the 0.2x ED50 bars indicate how many fold-lower gene expression was at the indicated dose as compared to the 1x ED50 dose. Thus, Cyp2b10 levels were 5.4-5.6 fold lower at 0.2x ED50 than at 1x ED50 (mature and primary RNA, respectively); and corresponding values were 8.5 and 3.4 fold higher for Cyp2c55 (mature and primary RNA, respectively). *: p < 0.05, **: p < 0.01 and ***: p < 0.001, pair-wise t-test for comparisons between each treatment and the vehicle control (first bar).

**Fig. S2.**
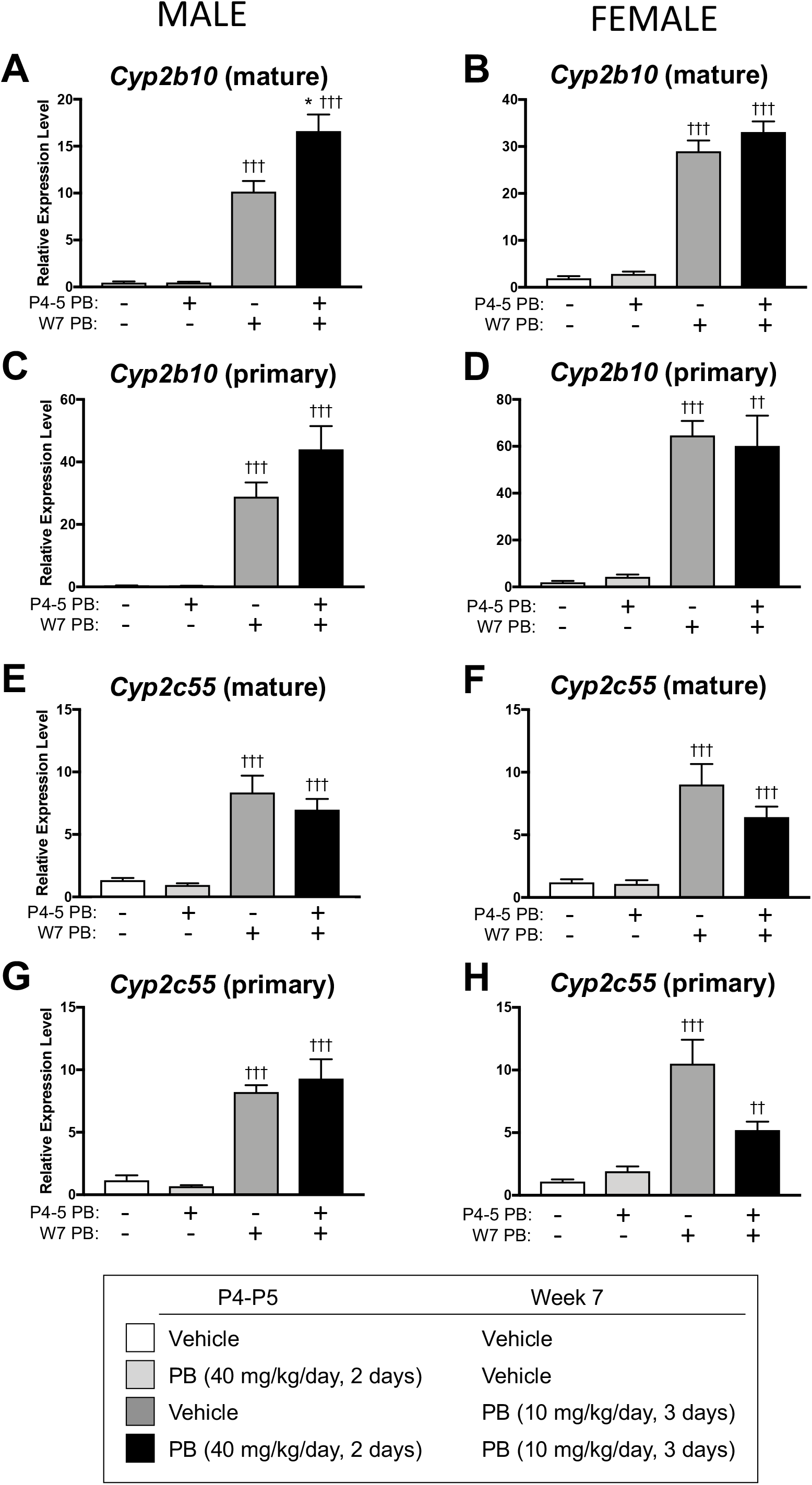
Impact of neonatal phenobarbital on PND4 and PND5 (40 mg/kg on each day) with re-challenge in week 7 (10 mg/kg daily for 3 consecutive days) (Study design D). Mice were euthanized 3 h after the last phenobarbital injection. Shown are total liver RNA expression levels determined by RT-qPCR for the mature and primary transcripts of Cyp2b10 and Cyp2c55, mean and SEM error bar for n = 4-8 livers per group in both male and female mouse livers. Pair-wise t-tests comparing: 1) neonatal phenobarbital vs vehicle control (bar 2 vs bar 1, and bar 4 vs bar 3: *: p < 0.05, **: p < 0.01, ***: p < 0.001; and 2) week 7 phenobarbital vs vehicle control (bar 3 vs bar 1, and bar 4 vs bar 2: †: p < 0.05, ††: p < 0.01, †††: p < 0.001)

**Fig. S3.**
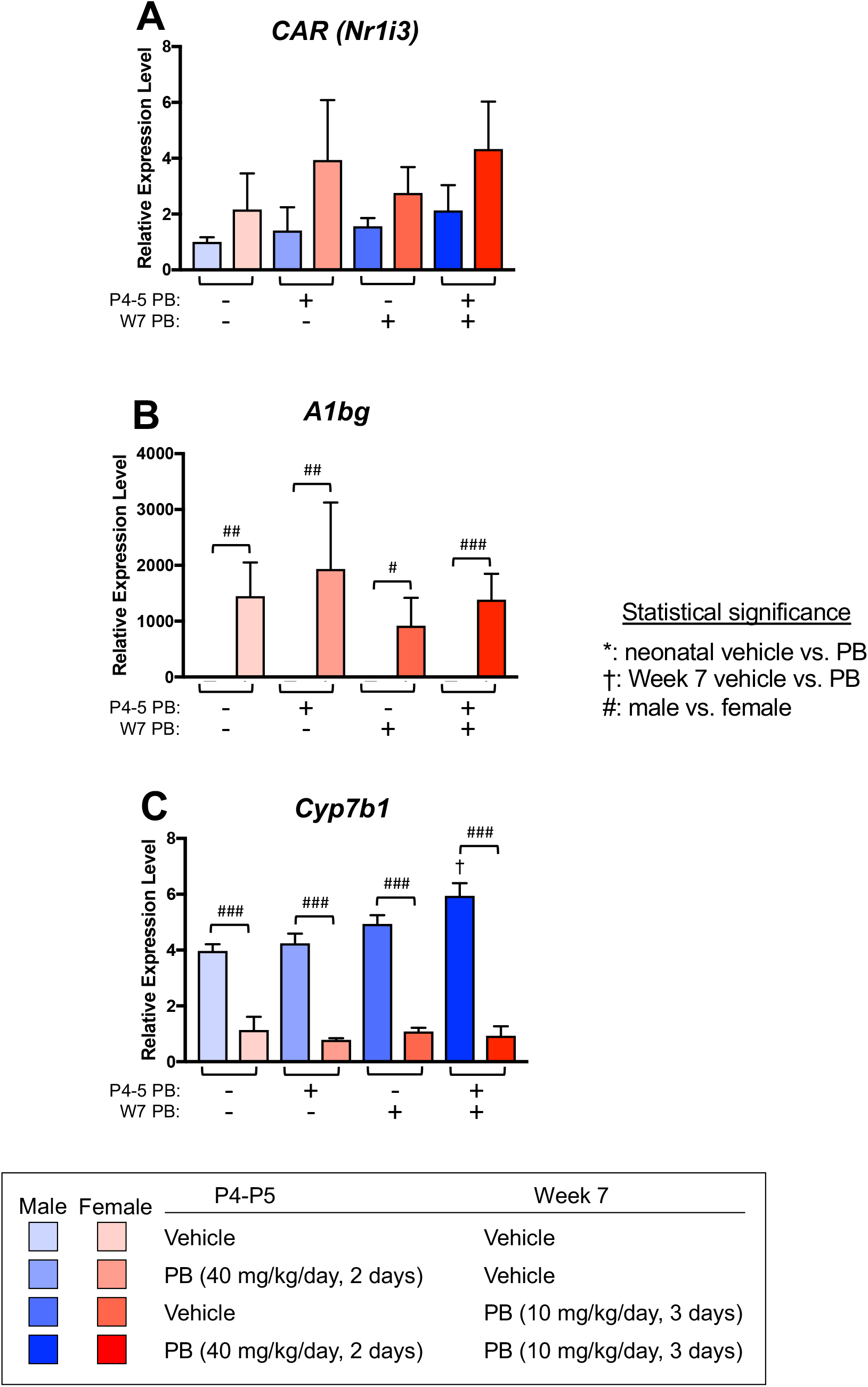
Impact of neonatal phenobarbital and adult phenobarbital rechallenge on expression of CAR and two sex-specific genes. Male and female mouse pups were injected either with vehicle or with phenobarbital at 40 mg/kg/day of phenobarbital on PND4 and P5, and later in week 7, were re-challenged with either vehicle or 10 mg/kg/day of phenobarbital for 3 consecutive days. Mice were euthanized at 3 h after the last phenobarbital injection. Shown are total liver RNA expression levels determined by RT-qPCR and normalized to the vehicle control group for CAR, for the female-specific A1bg, and for the male-specific Cyp7b1. Data shown are mean and SEM error bar for each group, for n = 3-6. Pair-wise t-tests comparing: 1) neonatal phenobarbital vs vehicle control: nothing significant at p < 0.05; 2) week 7 phenobarbital vs vehicle control (bar 3 vs bar 1, and bar 4 vs bar 2: †: p < 0.05); and 3) males vs females for the indicated 4 sets of comparisons, where #: p < 0.05, ##: p < 0.01 and ###: p < 0.001.

**Table S1.**
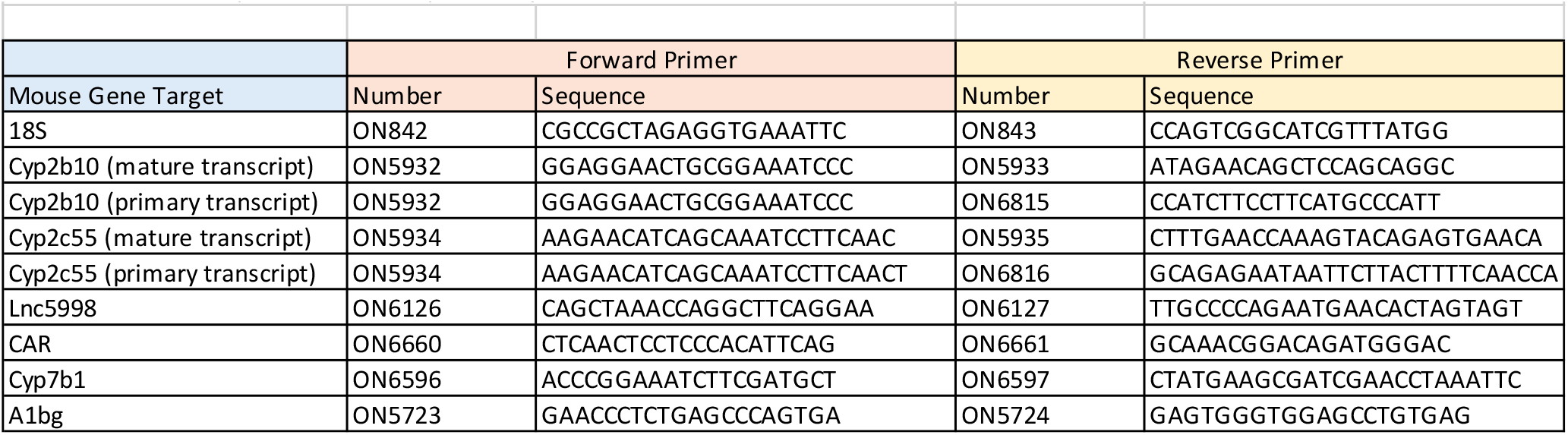
Primer sequences used for qPCR analysis.

